# Differential effects of conventional transcranial direct current stimulation (tDCS) and high-definition transcranial direct current stimulation (HD-tDCS) of the cerebellum on offset analgesia

**DOI:** 10.1101/2024.10.03.616281

**Authors:** Niamh O’Connor, Hannah Ashe, Max Wragan, Ruairí O’Flaherty, Eoin Deevy-Gray, Alice G. Witney

## Abstract

**Background:** Offset analgesia (OA) describes the large decrease in perceived pain in response to a minor decrease in applied painful thermal stimulus. Here non-invasive brain stimulation (NIBS) is used to modulate the cerebellum, since the cerebellum is known to signal sensory prediction errors and is implicated in pain processing.

**Methods:** An OA protocol individualized to heat pain threshold (HPT) was applied via TSA-II (Medoc, Israel). NIBS interventions were applied prior to OA. Cathodal cerebellar transcranial direct current stimulation (tDCS) and high-definition (4X1) transcranial direct current stimulation (HD-tDCS) were applied to 46 healthy participants within a sham controlled repeated measures design to examine whether diffuse or focal stimulation differentially modulates OA.

**Results:** OA induced hypoalgesia was robust, with 90% of responses showing a drop in perceived pain (δVAS) > 10 following the 1°C fall in temperature. This OA response was augmented following a protocol with sham and focal cathodal cerebellar stimulation on four OA parameters (OA _latency_, VAS _minimum_, VAS _mean_ and VAS _2nd max_) relative to pre-stimulation. This effect was differential to the protocol with sham and conventional tDCS where two OA metrics altered (OA _duration_, VAS _2nd max_).

**Conclusion:** OA enhancement via cathodal cerebellar NIBS may involve both a placebo effect and sustaining a noxious sensory prediction error. Understanding how the cerebellum is involved in OA could enhance therapies for pain patients.

## Introduction

Offset analgesia (OA) is a pain modulation phenomenon where an individual experiences a disproportionately large decrease in pain sensation in response to a small decrease in experimentally applied painful thermal stimulus [15; 17]. The transient unexpected decrease in applied pain leads to the experience of analgesia and hypoalgesia in 70-90% of healthy participants[17]. This OA response is thought to be due endogenous pain regulation systems with research demonstrating that abnormal or absent OA responses in chronic pain patients [1; 35; 37; 50]. Therefore, the OA protocol has potential to provide valuable insights into the function of endogenous pain regulation systems. These pathways are obvious therapeutic targets, so there is clinical utility in understanding the mechanisms underlying the strong analgesic effect that occurs following the unexpected drop in nociceptive thermal stimuli.

The cerebellum is known to contribute to pain modulation, but the precise role in pain processing has not been established [22; 32; 47]. The cerebellum has been widely studied in computational neuroscience due to the potential to map its geometric structure to theoretical models of learning, where a mismatch between predicted and actual sensory signals can provide an error correction mechanism that drives adaptation[30]. Further this learning process can be mapped to the cellular architecture of the cerebellum[19]. The cerebellar cortex has a modular organization with repeated microzones with a repeated simple neuronal arrangement [41; 42]. In each microzone, climbing fibres from the inferior olive project to Purkinje cells which then output to cerebellar and vestibular nuclei. Frameworks for learning suggest that synaptic plasticity, particularly long-term depression (LTD) drive error correction and motor learning. These ideas have led to the concept of the cerebellum involvement in the learning and updating an ‘internal model’, with this framework having broader applications than motor control[57]. Traditionally these frameworks are driven by self-generated movement, via efferent copies, predicting outcome. Applied to pain, this approach could explain the development and persistence of pain avoidance during movement. However, it may be that there is not a purely motor basis for such predictions[8]. The cerebellum’s role in pain processing may have a broader scope in the integration of nociception beyond the link with the motor system[44]. Offset analgesia could be explained via a predictive model, with the mismatch between actual and expected nociceptive stimuli leading to a re-calibration of the pain perceived via the same error correction mechanism.

Non-invasive brain stimulation (NIBS) is a means of modulating neuronal activity and has been found effective for pain modulation[3; 11; 12; 20; 23; 31]. Currently, the greatest efficacy for pain relief is associated with high frequency repetitive transcranial magnetic stimulation (rTMS) of primary motor cortex (M1) and anodal transcranial direct current stimulation (tDCS) of primary motor cortex (M1) [13; 21; 25]. Anodal M1 HD-tDCS has been found to modulate pain thresholds [55] with increased focality. These forms of M1 NIBS lead to increased corticomotoneuronal and corticocortical excitability with long term potentiation (LTP) like mechanisms proposed to underlie the synaptic plasticity. In contrast, cathodal tDCS is thought to reduce corticomotoneuronal excitability and induce LTD like changes. Within experimental studies of the cerebellum, both excitatory and inhibitory cerebellar stimulation have been shown to lead to modulation of pain thresholds in animal studies, dependent on the region stimulated[43; 46]. Further, the higher folding, increased neuronal density and wider diversity of cell types of the cerebellum relative to M1 add complexity to designing appropriate cerebellar NIBS protocols and may necessitate focal stimulation[53]. Some evidence has shown that cathodal tDCS may inhibit motor adaptation [18; 56]with the proposal that this can be explained via an inhibition of the error correction learning. If applied to pain, inhibition of updating the mismatch between actual and expected nociception could be predicted to sustain OA induced analgesia. Therefore, this study applies inhibitory cathodal NIBS to the cerebellum, comparing diffuse tDCS with the more focal HD-tDCS.

## Material and methods

The study protocol was approved by the Faculty of Health Science Research Ethics Committee, Trinity College Dublin, Ireland and conducted in accordance with the Declaration of Helsinki (2013) with pre-registration on the Open Science Framework. Participant eligibility was evaluated with a medical questionnaire, which excluded participants with neurological and psychiatric conditions. Power calculations determined that at least 44 participants were required. 46 participants (22 females, mean age 22.35 years) completed both experiments in the study. A sham-controlled repeated measures design was used, with tDCS and HD-tDCS completed in two sessions, separated by at least one week, to ensure no overlap in NIBS effects between experimental sessions. The order of the two NIBS interventions was counter- balanced between participants. Sham stimulation enables placebo NIBS effects on OA to be examined. Sham stimulation occurred prior to active stimulation in all participants in each session to avoid after-effects of active neurostimulation.

### Offset analgesia protocol

A TSA-II Neurosensory Analyzer, Medoc, Israel [29] applied heat via the 16x16 mm peltier thermode to the participants right forearm, secured with Velcro©. A computerized visual analogue scale (CoVAS, Medoc, Israel) [28] continuously recorded participants’ real-time pain report at 9 Hz [60] with the displacement of a marker between two endpoints of ‘no pain’ and ‘most intense pain imaginable’ corresponding to values between 0 and 100. Training on CoVAS occurred prior to testing.

Heat pain threshold (HPT) were determined to individualize the subsequent OA protocol [2; 33; 45; 48; 49]. HPTs were determined by increasing the temperature of the thermode applied to the right forearm at a rate of 1.5°C/s from a baseline of 32-35°C until pain was reported. HPT trials were repeated three times, with 10 s rest intervals between trials.

The OA protocol comprised of an OA trial and a control trial, presented in random order. To ensure data accuracy, 60 s of rest followed data collection trials in addition to slight thermode repositioning on the forearm to prevent sensitization [59]. On an OA trial (Figure 1), three sequential temperatures (T1-3) were utilised: the initial stimulus temperature (T1) is the participant’s HPT, ramped from the baseline temperature representing normal skin temperature (32-35°C) at a rate of 1.5℃/s. T1 (HPT) was held constant for 7 - 10 seconds. The duration of T1 varied to decrease participants’ expectation of a temperature change at temperature 2 (T2). The T2 phase elevates the temperature a single degree to HPT + 1℃ for 5 s, and the Temperature 3 (T3) falls to HPT for 30 s before returning to baseline skin temperature at a rate of 6℃/s. (Figure 1). The transition from T2 to T3 is where OA is experienced, the characteristic disproportionate decrease in pain sensation relative to the associated temperature decrease. This OA protocol differs from some previous OA protocols [15; 59] because the T3 phase was extended to characterize the duration of OA, previous studies show ongoing OA at the termination of T3 [2; 15; 51].

**Figure 1:**
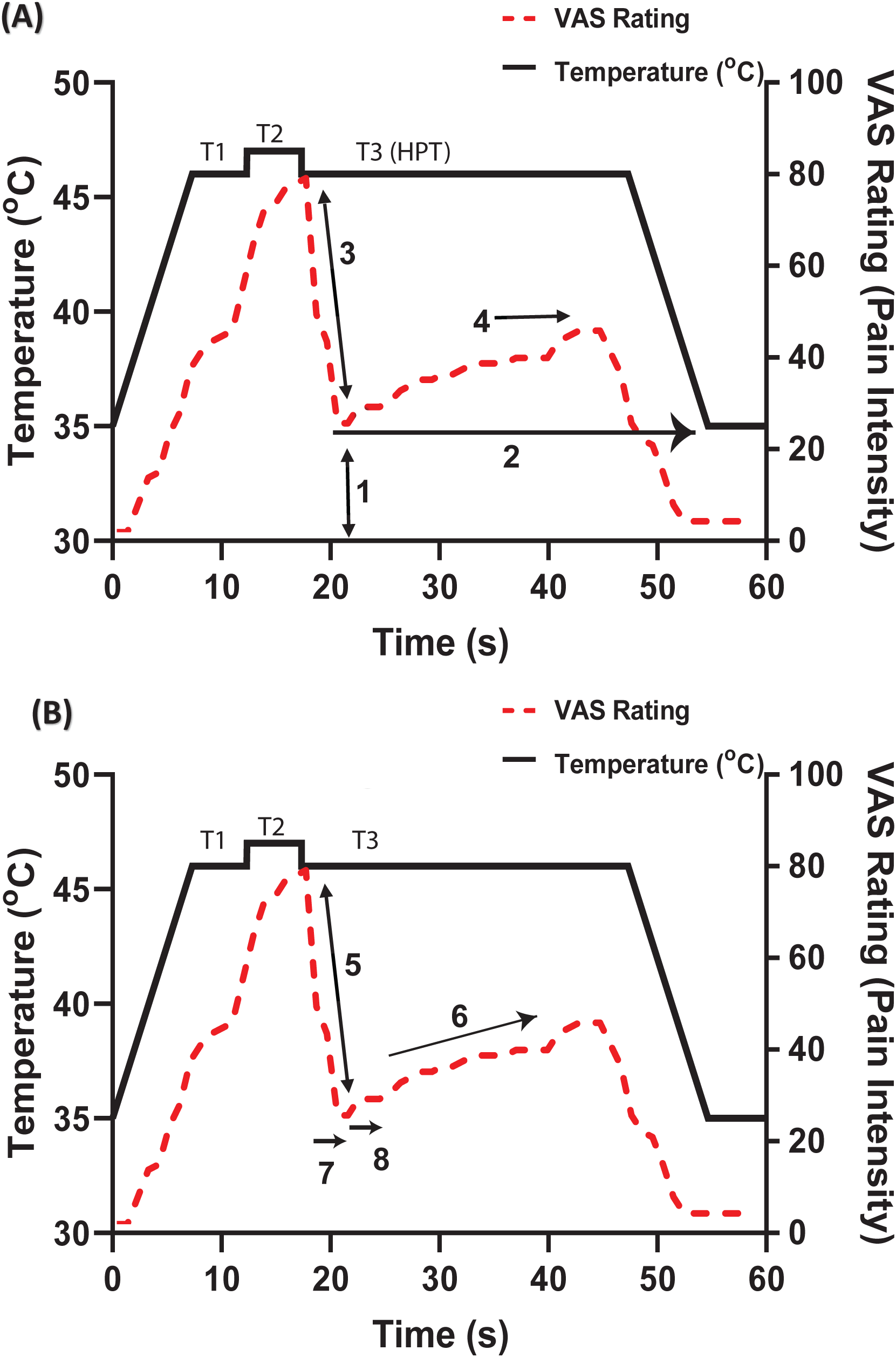
Offset analgesia protocol showing extracted OA metric definitions for magnitude (A) 1: VAS _min_, 2: VAS _mean_,3: δVAS _OA_ and temporal (B) 5: OA rate _down_, 6: OA _rate up_, 7: OA _latency_ and 8: 0A _duration_ of the OA response. See Table 1 for OA metric definitions.

**Table 1:** OA response metric definitions for magnitude and temporal features (see Figure 1 for graphical representation)

| Metric Label | Offset Analgesia Metric | Description |
| --- | --- | --- |
| 1 | VAS<sub>min</sub> | Lowest reported VAS in t3 |
| 2 | VAS<sub>mean</sub> | Mean VAS in t3 |
| 3 | δVAS<sub>OA</sub> | Magnitude of OA: the change in VAS at t2-t3 |
| 4 | VAS<sub>2nd Max</sub> | Maximum VAS in t3 following hypoalgesia |
| 5 | OA<sub>rate down</sub> | Rate of decrease in VAS (δVAS ) due to OA |
| 6 | OA<sub>rate up</sub> | Rate of increase from VAS<sub>min</sub> to VAS<sub>2nd Max</sub> |
| 7 | OA<sub>latency</sub> | Time taken for maximum VAS score at the start of t3 to drop to the OA nadir, VAS<sub>min</sub> (the OA induced hypoalgesia) |
| 8 | OA<sub>duration</sub> | Time taken for VAS<sub>min</sub> to rise again following hypoalgesia |

Control trials in the OA protocol consisted of the same T1 temperature as the OA trials (HPT), similarly ramped from baseline skin temperature at a rate of 1.5℃/s. However, in contrast, T1 was sustained for 30s, before ramping down to baseline skin temperature at rate of 6℃/s.

### Transcranial direct current stimulation montages

Conventional cerebellar tDCS was applied using a neuroConn DC stimulator (NeuroConn GmBH, Germany) with sponge electrodes (surface area 25 cm^2^) soaked in 0.9% saline. The stimulating electrode was positioned at the mid-point between the inion and the maximum convexity of the right ear with the return electrode on the right buccinator[16; 18; 24].

Cerebellar HD-tDCS was applied using the neuroConn DC stimulator connected to a Soterix 4x1 HD-tDCS adaptor (Model 4x1-C2; Soterix Medical Inc., New York, USA). A 4X1 electrode montage using sintered Ag/AgCl electrodes (EasyCap, GmbH, Germany) filled with conductive gel (Electro-Gel, Brainmaster Inc., USA) were secured with an electrode cap (EasyCap, GmbH, Germany). The central electrode was placed at the mid-point of the inion and the maximum convexity of the right ear, with the four return electrodes spaced 3 cm radially around the central electrode. Impedance values were measured for all electrodes prior to usage to ensure good contact quality [54; 55]. Active NIBS applied current of 2mA of tDCS for 20 minutes for both NIBS protocols. Sham stimulation applied 2mA for 30s to elicit a tingling sensation, with no current applied in remainder of the 20 minute protocol [14; 53].

Current flow modelling is critical to NIBS protocol design[10; 39]. The cerebellar tDCS and HD-tDCS protocols were simulated (HD-Explore Version 2.1, Soterix Medical, New York, US) to characterise the current distribution with the given electrode montage and current level of 2mA as applied experimentally. All modelling was performed on a 93 standard adult male head model sourced from the Soterix model library. Electrode positioning was based on the 10/20 EEG system. For conventional tDCS modelling, O10, PO10, EX2, and EX6 represented the cathode sponge electrode and EX14, EX16, and EX18 represented the anodal sponge electrode (Figure 2a). For HD-tDCS modelling, PO10 represented the cathode and PO4, P10, EXZ, and EX4 represented the anodes (Figure 2b).

**Figure 2:**
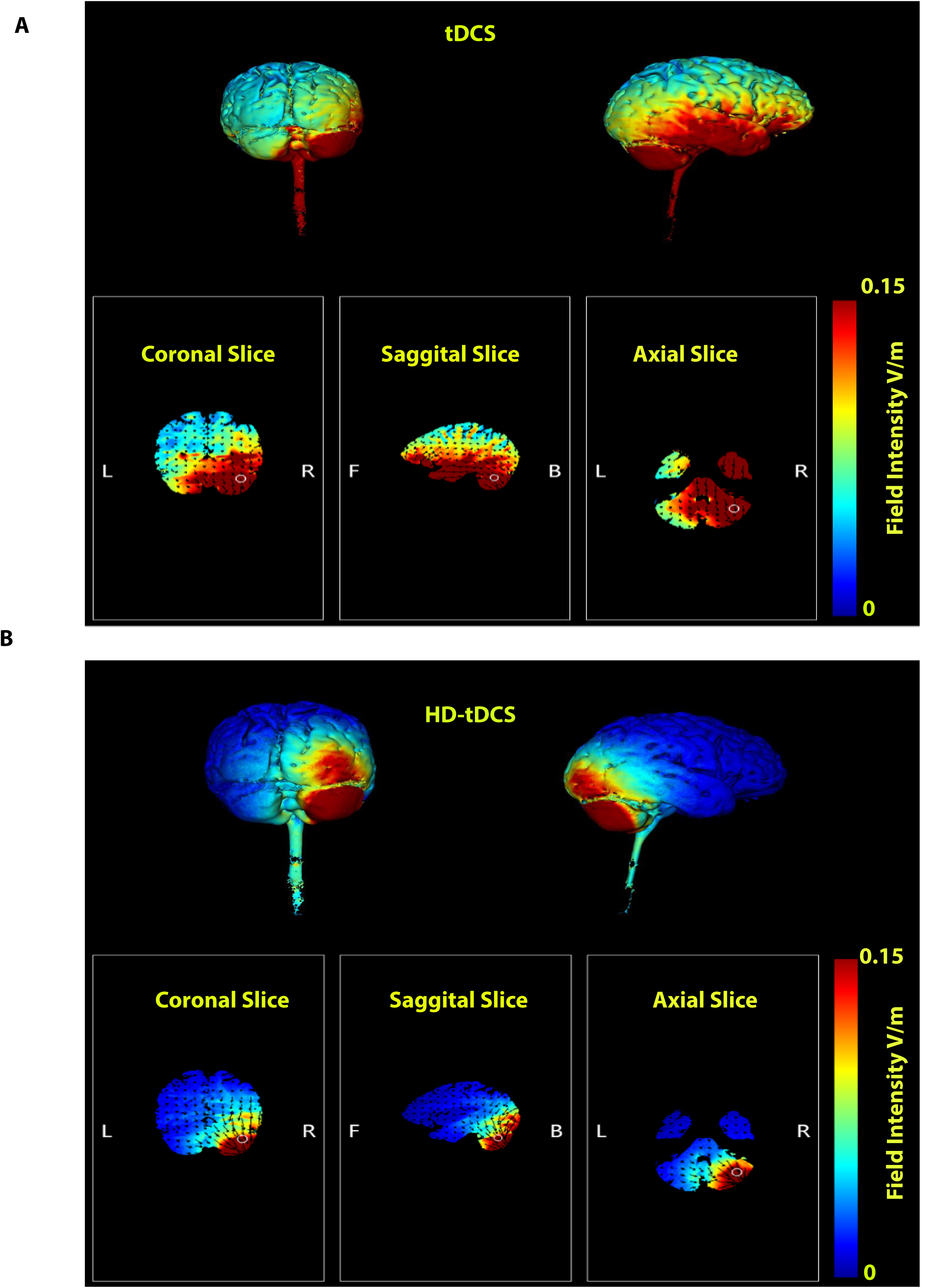
Current flow modelling: 3D representation of current flow (upper panel), current flow through coronal, sagittal and axial slices (lower panel) A) Cerebellar tDCS montage: High intensity in the cerebellum, temporal lobe, frontal lobe and brainstem. B) Cerebellar HD-tDCS montage restricted current to the right cerebellar hemisphere and the posterior occipital lobe, with peak field intensity in the posterior cerebellum.

### OA and control trial response characterization

OA and Control response metrics were extracted using a custom written MATLAB script (Version 9.0, The MathWorks Inc., MA, USA). Control trial metrics were mean (VAS _mean_), minimum (VAS _min_), maximum (VAS_max_) and standard deviation (VAS_sd_) of the VAS score while HPT was applied over a 30s period (T1).

CoVAS profiles were averaged dependent on neurostimulation (pre-stimulation, sham, active stimulation) from the onset T3 for 22 s to qualitatively compare the impact of NIBS on the OA response, which is defined as the VAS ratings made during T3. The OA response metrics used in the study (Table 1, Figure 1) included both magnitude (Figure 1a) and temporal (Figure 1b) features with metrics selected to be consistent with previous studies, VAS _min_ [2; 6; 35], δVAS [1; 6; 35] and VAS _mean_ [6]. Further, in this study, a second maximum VAS was included since the OA protocol extended the T3 phase. Some OA metrics could not be extracted on all trials due to features of the OA response, for example there could be no VAS _2nd max_ or OA _rate up_ metric if the OA continued throughout T3.

### Statistical analysis

Statistical analysis was performed in SPSS (IBM SPSS Statistics, Version 27; IBM Corp). Descriptive statistics were collected for all OA metrics across conditions (pre-stimulation OA; sham stimulation OA; active stimulation OA). Within day ICC on OA metrics are consistently high in previous research [36] and pilot work in the laboratory. Between day ICC comparing OA metrics at pre-stimulation baseline on each session were moderate or low. Therefore, the tDCS and HD-tDCS experiments were considered separately via two 3X1 repeated measures ANOVA for both control and OA trial metrics. Mauchly’s test of sphericity was performed, and when sphericity could not be assumed the Huynh-Feldt estimate was applied. Post-hoc pairwise comparisons were made with Bonferroni corrections for multiple comparisons.

## Results

On control trials where T1 was held at HPT for 30s, the VAS _mean_ was 30.07 (SD 18.95), VAS_max_ of 42.31 (SD 22.41), VAS _min_15.96 (SD 18.67) and VAS _sd_ was 28.11 (SD 9.06). As expected, there were no consistent periods of analgesia observed in this VAS response to sustained HPT.

In contrast, the OA trials successfully consistently induced analgesia, with only 3% of trials not showing a decrease in VAS at the T2-T3 transition. 90% of trials showed a fall greater than 10 VAS, with most trials eliciting a much larger decrease in pain perception.

The fall of temperature in T3 to HPT gave hypoalgesia with a minimum VAS (VAS _min_) at the nadir of the OA response of 12.71 (SD 15.50). The mean δVAS in the current study was slightly lower at 38.13 (SD, 22.39) relative to scores found by previous studies [1; 33; 34]. The T3 VAS _mean_ 30.35 (SD 23.61) for T3 phase was lower than studies that do not individualize OA protocols to HPTs [6]. The OA _rate down_ at the T2-T3 transition a mean of 8.45 VAS/s (SD 8.45) was much steeper than OA _rate up_, during T3, (1.85 VAS/s, SD 1.71), revealing the rapidity of OA at the transition and the sustained hypoalgesia that demonstrates the necessity of extending T3 in the protocol. Taken together the protocol used in this study robustly generated OA and provided baseline values consistent with earlier studies in the area [1; 27; 33].

### Modelling cerebellar NIBS

Current flow in the conventional tDCS (Figure 2a) is more diffuse than HD-tDCS (Figure 2b). Highest flow intensities with tDCS are near the active electrode, at the cerebellar cortical level in the posterior lobe. Medial structures also receive strong current and spread to the temporal lobe and frontal lobe. Current reached the upper and lower brainstem. For HD- tDCS, current flow was more restricted, with high field intensities in the posterior and anterior cerebellar lobe and posterior aspects of the temporal lobe. Very little current reaches the contralateral cerebellar hemisphere or brainstem.

### The effect of cerebellar NIBS on the OA protocol

There were no significant differences in the control trial metrics (VAS _mean_, VAS _min_, VAS_max_ and VAS_sd_) dependent on stimulation for either tDCS or HD-tDCS. However, there was a differential impact of NIBS on the OA response (Figure 3).

**Figure 3:**
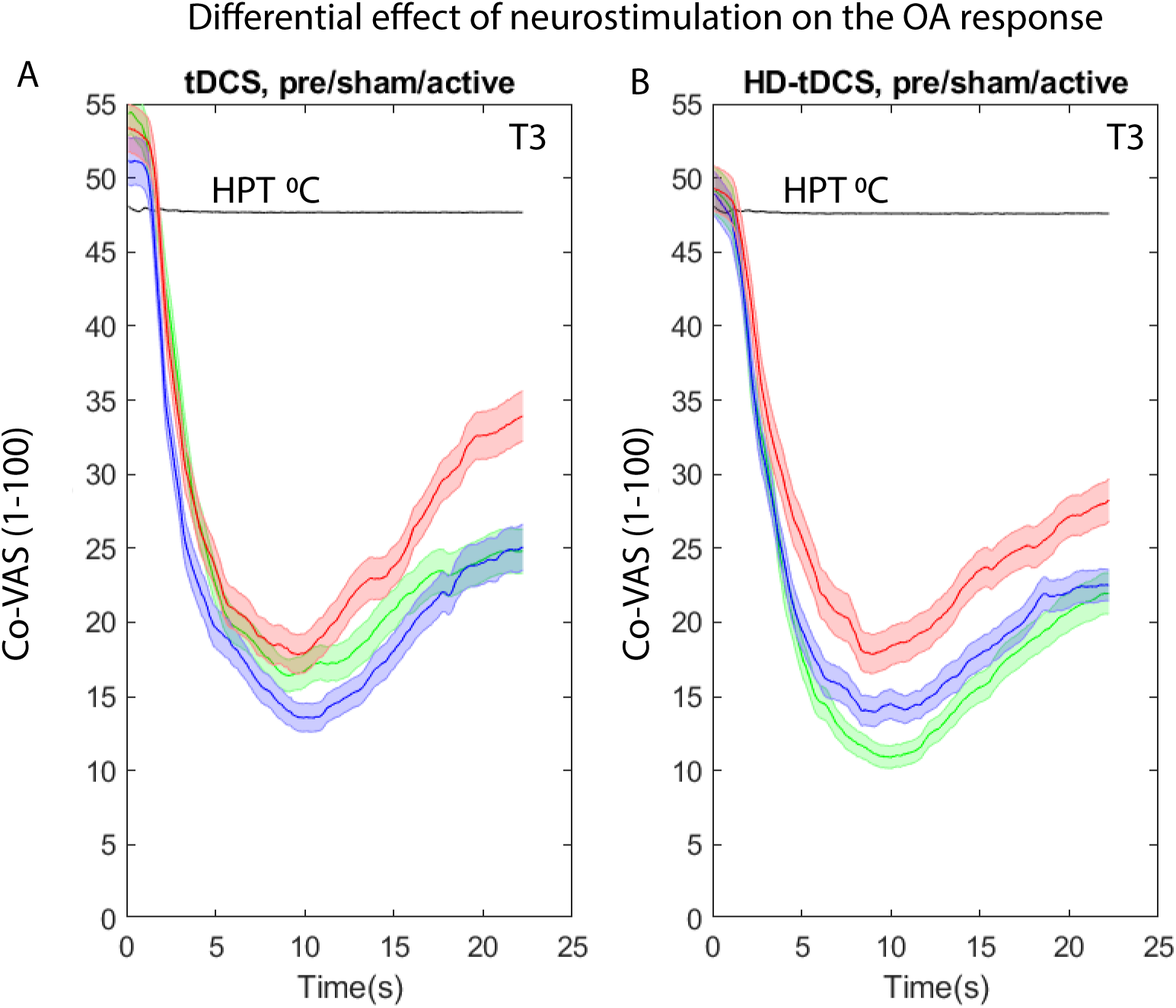
Average of OA response during T3 period dependent on cerebellar NIBS stimulation for a) tDCS protocol and b) HD-tDCS protocol. Pre-stimulation (red); Sham- stimulation (blue); Active-stimulation (green). Shading represents standard error of the mean.

The average impact of active conventional cerebellar tDCS stimulation relative to pre- stimulation and sham can be seen in Figure (3a). Mean values of the OA pain metrics are found in table 2.

**Table 2:** OA pain profile metric means.

| stimulation | condition | VASmin | VASmean | δVAS | VAS<br>(2<sup>nd</sup><br>max) | OA<br>Rate<br>down<br>(VAS/s) | OA<br>Rate<br>Up<br>(VAS/s) | OA<br>Latency<br>(s) | OA<br>duration<br>(s) |
| --- | --- | --- | --- | --- | --- | --- | --- | --- | --- |
| tDCS | Pre | 14.74<br>(16.67) | 26.77<br>(20.62) | 32.70<br>(20.26) | 27.79<br>(18.98) | 6.59<br>(4.75) | 1.41<br>(1.40) | 6.09<br>(3.31) | 8.11<br>(6.75) |
|  | Sham | 10.08<br>(11.71) | 23.51<br>(17.62) | 40.19<br>(23.76) | 29.52<br>(25.33) | 9.80<br>(10.07) | 2.10<br>(1.78) | 5.93<br>(3.34) | 10.48<br>(8.59) |
|  | Active | 14.29<br>(16.37) | 25.23<br>(18.17) | 40.06<br>(22.63) | 29.75*<br>(23.80) | 9.08<br>(9.81) | 1.96<br>(1.45) | 6.20<br>(3.50) | 11.17*<br>(7.97) |
| HD-tDCS | Pre | 16.17<br>(20.25) | 28.77<br>(19.59) | 37.80<br>(22.40) | 37.35<br>(26.94) | 8.04<br>(7.65) | 1.87<br>(1.76) | 6.23<br>(3.35) | 9.80<br>(6.75) |
|  | Sham | 11.35<br>(14.56) | 21.49<br>(13.49) | 38.02<br>(23.51) | 27.79<br>(18.99) | 9.80<br>(10.08) | 1.81<br>(1.87) | 5.53<br>(2.97) | 8.90<br>(6.84) |
|  | Active | 9.79*<br>(11.62) | 20.91*<br>(15.19) | 39.75<br>(21.79) | 26.02*<br>(21.7) | 8.31*<br>(5.37) | 1.96<br>(1.90) | 5.69<br>(2.92) | 10.62<br>(7.69) |

### The effect of sham conventional tDCS and cerebellar cathodal tDCS on OA trials

The conventional tDCS protocol led to two significant changes in OA metrics with moderate effect sizes. The alteration in OA _duration_ was found to be significant F(2,86)=4.35, p =.037, partial η^2^ = .07. Post-hoc pairwise comparisons with a Bonferroni adjustment indicated there was no significance difference between OA _duration_ at baseline and following sham stimulation, p=.18. However, OA _duration_ was significantly longer after active tDCS stimulation (11.17 s, SD 7.97) relative to pre-stimulation (8.11 s, SD 6.75), p=.048. Interestingly, there was also no significant difference between OA _duration_ after sham and after active neurostimulation, p=1. OA _duration_ represents the length of maximal OA; from the T2-T3 transition to the point at which pain rating begins to rise again (Figure 1b, Table 1, metric 8).

Additionally, there was an effect of stimulation of VAS _2nd max_, F(2,90)=5.40,p=.006, partial η^2^ = .11. Post-hoc pairwise comparisons with a Bonferroni adjustment showed there was a significant difference between mean VAS _2nd max_ at baseline (37.35, SD 26.94) and following sham stimulation, (29.75, SD 25.32) p=.009 additionally VAS _2nd max_ was significantly lower after active tDCS stimulation, (29.52, SD 25.33), p=.028. However, the reduction in VAS _2nd max_ also occurred after sham stimulation, with no significant difference between VAS _2nd max_ and after active stimulation, p=1, suggestive of a placebo effect. Statistical analysis revealed there was no effect of cerebellar tDCS neurostimulation on δVAS, VAS _min_, OA _rate down_, VAS _mean_, OA _rate up_ or OA l_atency_ metrics of the OA response.

### The effect of sham HD-tDCS and cathodal cerebellar HD-tDCS on OA trials

Qualitatively, the effect of active cathodal cerebellar HD-tDCS relative to sham is to enhance the OA effect (Figure 3b). Statistical analysis revealed differences on four OA metrics with medium effect sizes. The effect of neurostimulation on δVAS, that is the drop in VAS at the T2-T3 transition was found to be significant F(1.64,76.09)=3.72, p =.036, partial η^2^ = .08. Post-hoc pairwise comparisons with a Bonferroni adjustment indicated there was no significance difference between mean δVAS at baseline (32.70, SD 22.63) and following sham stimulation, (38.02, SD 23.51), p=.46. However, δVAS was significantly greater after active stimulation, (39.80, SD=21.79), p=.007. There was also no significant difference between δVAS after sham and after active neurostimulation, p=.96.

The effect of neurostimulation on VAS _rate down_, that is the rate of OA-induced hypoalgesia, was found to be significant F(1.70,71.18)=3.36, p =.048, partial ^2^ = .07. Post-hoc pairwise comparisons with a Bonferroni adjustment indicated there was no significance difference between mean VAS _rate down_ at baseline (6.59 VAS/s, SD 4.75) and following sham stimulation, (8.91 VAS/s, SD=8.38), p=.55. However, VAS _rate down_ was significantly faster after active tDCS stimulation, (8.31 VAS/s, SD 5.37) p=.021. There was also no significant difference between VAS _min_ after sham and after active neurostimulation, p=.86.

The effect of neurostimulation on VAS _min_, the VAS at the nadir of the OA response was found to be significant F(2,90)=4.96, p =.01, partial η^2^ = .1. Post-hoc pairwise comparisons with a Bonferroni adjustment indicated there was no significance difference between mean VAS_min_ at baseline (16.17, SD=20.25) and following sham stimulation, (10.08, SD 11.62), p=.12. However, mean VAS_min_ was significantly reduced after active tDCS stimulation, (9.79, SD 11.62), p=.018. There was also no significant difference between VAS_min_ after sham and after active neurostimulation, p=1.

There was an effect of stimulation of VAS _mean_, F(2,90)=4.91, p =.009, partial η^2^= .98. Post- hoc pairwise comparisons with a Bonferroni adjustment indicated there was no significance difference between mean VAS_mean_ at baseline (32.04, SD 23.81) and following sham stimulation, p=.055. However, VAS _mean_ was significantly lower after active tDCS stimulation, (26.02, SD 21.75) p=.023. There was also no significant difference between VAS _mean_ after sham (27.79, SD 18.99), and after active neurostimulation, p=1.

The effect of neurostimulation on VAS_2nd max_ was found to be significant F(2,90)=3.56, p =.033, partial ^2^ = .07 indicating pain does not rise as much even though OA is reduced relative to the start of T3. Post-hoc pairwise comparisons with a Bonferroni adjustment indicated there was no significance difference between mean VAS_2nd max_ at baseline (32.04, SD 23.81) and following sham stimulation, (27.29, SD 18.99), p=.19. However, VAS_2nd max_ was significantly reduced after active focal tDCS stimulation, (26.02, SD 21.75), p=.038. There was also no significant difference between VAS_2nd max_ after sham and after active neurostimulation, p=1.

There were no effect of the NIBS protocol on OA _duration_, OA _latency_ or OA _rate up._

## Discussion

Offset analgesia, the disproportionate decrease in perceived pain that follows a slight decrease in applied nociceptive stimuli, is a robust experimental pain protocol[15]. One previously untested explanation for offset analgesia is that it is experienced due to sensory error signaling in the cerebellum. Error signaling in the cerebellum drives sensorimotor adaptation and motor learning in altering environments[57]. If this concept were applied to pain, a reduction in adaptation might prolong the analgesia evoked by the transient sensory mismatch in an OA trial. To address this, the study applied inhibitory NIBS to the cerebellum, to examine whether cathodal tDCS may reduce adaptation to the sensory mismatch implicit within the OA protocol and thus sustain analgesia. This prediction of the study was supported with focal NIBS, yet also it was further revealed that placebo effects could contribute to modulation of OA.

The study first designed an individualized offset analgesia protocol based on prior studies[17]. Consistent with earlier OA studies [15], the protocol provided a robust methodology for inducing hypoalgesia, with participants consistently experiencing a profound reduction in the magnitude of pain experienced during the protocol. One aim of this study was quantifying the OA temporal pain profile with metrics that could characterize both magnitude and duration of the experienced pain. The increased duration of an OA trial in this study protocol is an important modification to earlier protocols as it assists differentiation and characterization of OA pain profiles; the essential basis for determining the effects of NIBS on OA.

Focal, HD-tDCS was found to augment the effects of OA pain profiles in four of the metrics with medium effect sizes with conventional cerebellar tDCS augmenting the OA pain profile on two metrics relative to pre-stimulation. However, in this study the interpretation of NIBS effects on the OA response is likely to summative to the occurrence of repeated OA protocols and perhaps also to sham NIBS providing a placebo effect. Qualitatively there appears to be a trend towards increased OA after sham (Figure 3), and one metric in the conventional cerebellar tDCS experiment showed a significant difference post sham, suggestive that a placebo effect is present. However, no other OA metrics post-sham reach significance in either experiment. Therefore, it is possible that the effect of repeated OA with sham NIBS primes the OA response, with subsequent active NIBS intervention augmenting the effect, producing the overall significant changes to the OA pain profiles. Previous studies have investigated the impact of repeated OA, with mixed findings[5; 17]. Importantly, the differential impact of the two active NIBS interventions on OA is observable both qualitatively and statistically, indicating that interpretation must include consideration of current flow and the focality of current applied and not be solely a placebo effect.

Analgesic enhancement in OA can be explained by several different pathways, and cerebellar stimulation via NIBS can contribute to this enhancement. The exact mechanisms of NIBS on the cerebellum are less well established than cortical regions [53], partly due to the increased diversity of cell types in cerebellar cortex, though it is thought to modulate Purkinje cells. There are multiple possible consequences to Purkinje cell modulation, since these are the sole output cells from cerebellar cortex. First, cathodal cerebellar tDCS may decrease inhibitory effects of Purkinje cells and so increases deep cerebellar nuclei activity, to indirectly reduce inhibition of M1 [13]. Numerous studies link analgesia with primary motor cortex excitability and the cerebellum has a known role in modulating corticomotoneuronal activity[38]. This pathway forms the cerebello-thalamo-cortical pathway, a major output pathway of the cerebellum. Alternatively, increased caudate putamen activity is associated with cathodal and anodal cerebellar tDCS [25]. The caudate is a known region for pain suppression [58] and is critically involved in the reward pathways. The hypoalgesia experienced after the transient decrease in supra-threshold pain, that is characteristic of the OA protocol, is known to activate reward pathways. Additionally, cerebellar stimulation may impact on OA via connectivity with known pain modulation pathways including the PAG and dorsolateral prefrontal cortex (DLPFC)[4].

This study showed in a healthy young population, cathodal cerebellar NIBS modulated the OA response relative to pre-stimulation consistent with disruption of cerebellar adaptation. This is notable since the OA response in a healthy population such as the one in this study can be subject to ceiling effects. Future work would apply the study protocol to an elderly and chronic pain patient population, both of whom have deficits in OA. However, studies in healthy, young adults are first necessary to ensure a robust protocol, minimizing individual differences that would impact on the OA response. The differences in effects due to focality of stimulation are perhaps unsurprising given that animal studies show both pro and anti- nociceptive effects of the cerebellum[7; 46] and due to the relatively small target regions [52]. This suggests that future work should use high-definition montages, alongside current flow modeling. Cerebellar NIBS protocols and electrode montages are developing [9; 26; 53] but there is increasing evidence that the choice of optimal electrode montage and ‘dosage’ is critical for an optimal effect [40].

Overall OA is a robustly observed form of hypoalgesia and better understanding of the underlying mechanism could open therapeutic options. This study supports further investigation of cerebellar involvement in OA.

## Author contributions

NOC and AW designed the experimental protocol with EDG involved in pilot testing; NOC, HA, ROF collected the experimental data, MW and AW collated the data, MW and AW performed statistical and data analysis. NOC, HA, ROF, EDG contributed to initial drafts. MW and AW wrote the manuscript.

